# Optimal frequency of perinatal retinal waves is essential for the precise wiring of visual axons in non-image forming nuclei

**DOI:** 10.1101/2022.07.27.501692

**Authors:** Santiago Negueruela, Cruz Morenilla-Palao, Salvador Sala, Macarena Herrera, Yaiza Coca, Maria Teresa López-Cascales, Patricia Ordoño, Danny Florez-Paz, Ana Gomis, Eloísa Herrera

**Affiliations:** Instituto de Neurociencias de Alicante (Consejo Superior de Investigaciones Científicas-Universidad Miguel Hernández, CSIC-UMH). Campus San Juan, Ave. Ramón y Cajal s/n, 03550 San Juan de Alicante (Spain)

**Keywords:** Axon Remodeling, retinal spontaneous activity, Kir2.1, visual targets, suprachiasmatic nucleus

## Abstract

The development of the visual system is an intricate and multi-step process involving the precise connection of retinal ganglion cell (RGC) axon terminals with their corresponding neurons in the visual nuclei of the brain. Upon reaching primary image-forming nuclei (IFN), such as the superior colliculus and the lateral geniculate nucleus, RGC axons undergo extensive arborization that refines over the first few postnatal weeks. The molecular mechanisms driving this activity-dependent remodeling process, which is influenced by spontaneous activity in the developing retina, are still not well understood. In this study, by manipulating the activity of RGCs in mice and analyzing their transcriptomic profiles before eye opening, we have identified gene programs involved in activity-dependent refinement. Furthermore, while RGC axons also target non-image forming nuclei (NIFN), the impact of spontaneous retinal activity on the development of these accessory nuclei, has not yet been elucidated. The analysis of visual terminals from mice with altered retinal activity revealed that spontaneous retinal waves occurring prior to visual experience also play a role in shaping the connectivity of the non-image forming circuit. Overall, these findings contribute to a deeper understanding of the mechanisms governing activity-dependent axon refinement during the establishment of the visual circuit.

## INTRODUCTION

Retinal ganglion cells (RGCs) are the cells in the eye that collect visual information from other neurons in the retina and send it to different nuclei in the brain including the superior colliculus (SC) and the lateral geniculate nucleus of the thalamus (LGN). These nuclei, also known as image-formin nuclei (IFN) are both essential for sight processing. While the SC directs visually-guided behaviors, the dLGN relays visual information to the cortex for a deeper conscious visual perception (Stryker and Schiller, 1975). Neighboring RGCs projecting to neighboring cells in the dLGN and the SC form a continuous topographic map that reflects the image projected in the retina (Rakic, 1976; Godement et al., 1984; Chalupa and Snider, 1998). Thus, in the IFN the spatial relationship among the RGCs in the retina is maintained as an orderly representation (map) of the visual space. In addition to form topographic maps in the dLGN and SC, RGC axons project in an eye-specific manner in most mammals. In mice, axons from a subpopulation of RGC located in the ventrotemporal retina project to the SC and dLGN in the same hemisphere (ipsilateral RGCs), while the rest of RGC axons cross the brain midline and project to the opposite side (contralateral RGCs). Ipsi and contralateral terminals first overlap at early postnatal stages in the dLGN but eventually segregate to finally rearrange into complementary non-overlapping domains in the mature visual system (Godement et al., 1984).

Both the formation of a raw retinotopic map and the targeting of ipsi and contralateral visual terminals at the SC and the thalamus are initially governed by guidance and adhesion molecules (Nakamoto et al., 2019; Su et al., 2021). In rodents, spontaneous waves of correlated activity triggered by starbust amacrine cells and transmitted to RGCs, refine and prepare the network to receive environmental stimuli when eyes open around two weeks after birth (Galli and Maffei, 1988; Wong et al., 1993; Demas et al., 2003; McLaughlin et al., 2003; Pak et al., 2004; Torborg and Feller, 2005; Torborg et al., 2005). However, the molecular mechanisms that shape the visual circuit by transducing retinal spontaneous activity in axon remodelling at the IFN remain largely unknown.

In addition to project to IFN, RGC axons project to NIFN such as the olivary pretectal nucleus (OPN), the nucleus of the optic tract (NOT) or the suprachiasmatic nucleus (SCN). The OPN and NOT mediate the pupillary reflex (Young and Lund, 1994) and saccadic movements (Kato et al., 1986) for image stabilization while the SCN regulates entrainment of the circadian clock, hormone rhythms and sleep cycles (Hattar et al., 2003; Yonehara et al., 2009; Noseda and Burstein, 2011; Dhande et al., 2013). Correlated retinal activity encodes positional information essential for the remodeling of RGC axons at the IFN but the impact of spontaneous retinal activity on the arborization of RGC terminals in the NIFN has not been investigated. NIFN receive inputs from the intrinsically photosensitive retinal ganglion cells (ipRGCs), a particular type of RGCs that responds directly to light and expresses melanopsin photopigment (OPN4) (Berson et al., 2002; Hattar et al., 2002; Do and Yau, 2010; Schmidt and Kofuji, 2011). NIFN work unconsciously to indirectly support vision without apparent need to transmit spatial information. Therefore, the organization of RGC axons in topographic maps and eye - specific projection patterns may not be as evident or even absent in NIFN.

Here, by generating a transgenic mouse line with altered retinal waves and analyzing the transcriptomic profile of their RGCs we retrieved molecular signatures that respond to proper spontaneous activity and are involved in the refinement of retinal axons. Moreover, the analysis of retinal terminals in the different visual nuclei of these mutant mice with altered retinal waves, demonstrated that proper frequency of perinatal retinal waves is essential for the establishment of accurate connectivity in non-image forming visual circuits.

## RESULTS

### Generation and characterization of mice ectopically expressing Kir2.1 in RGCs

To investigate the underlying activity-dependent genetic program that modifies axon remodeling at the visual targets and a potential function of spontaneous retinal waves in NIFN, we aimed to generate a mouse line with disrupted embryonic retinal activity by conditionally expressing the rectifying potassium channel Kir2.1 in RGCs. We first generated a line containing hKir2.1 fused to the fluorescence protein eYFP in a cre-dependent manner (Kir2.1YFP^flxStop^) and breed these mice with a Pou4f2-cre to drive gene expression in recently differentiated RGCs (Fuerst et al., 2012) obtaining double mutant mice Pou4f2-cre;Kir2.1YFP^fxStop^ (Pou4f2-Kir2.1) (**Fig. 1A**). The analysis of retinal sections from newborn Pou4f2-Kir2.1 mice confirmed stronger expression of hKir2.1eYFP in RGCs compared to the retinas of control littermates (Kir2.1YFP^flxStop^) (**Fig. 1B**).

**Figure 1.**
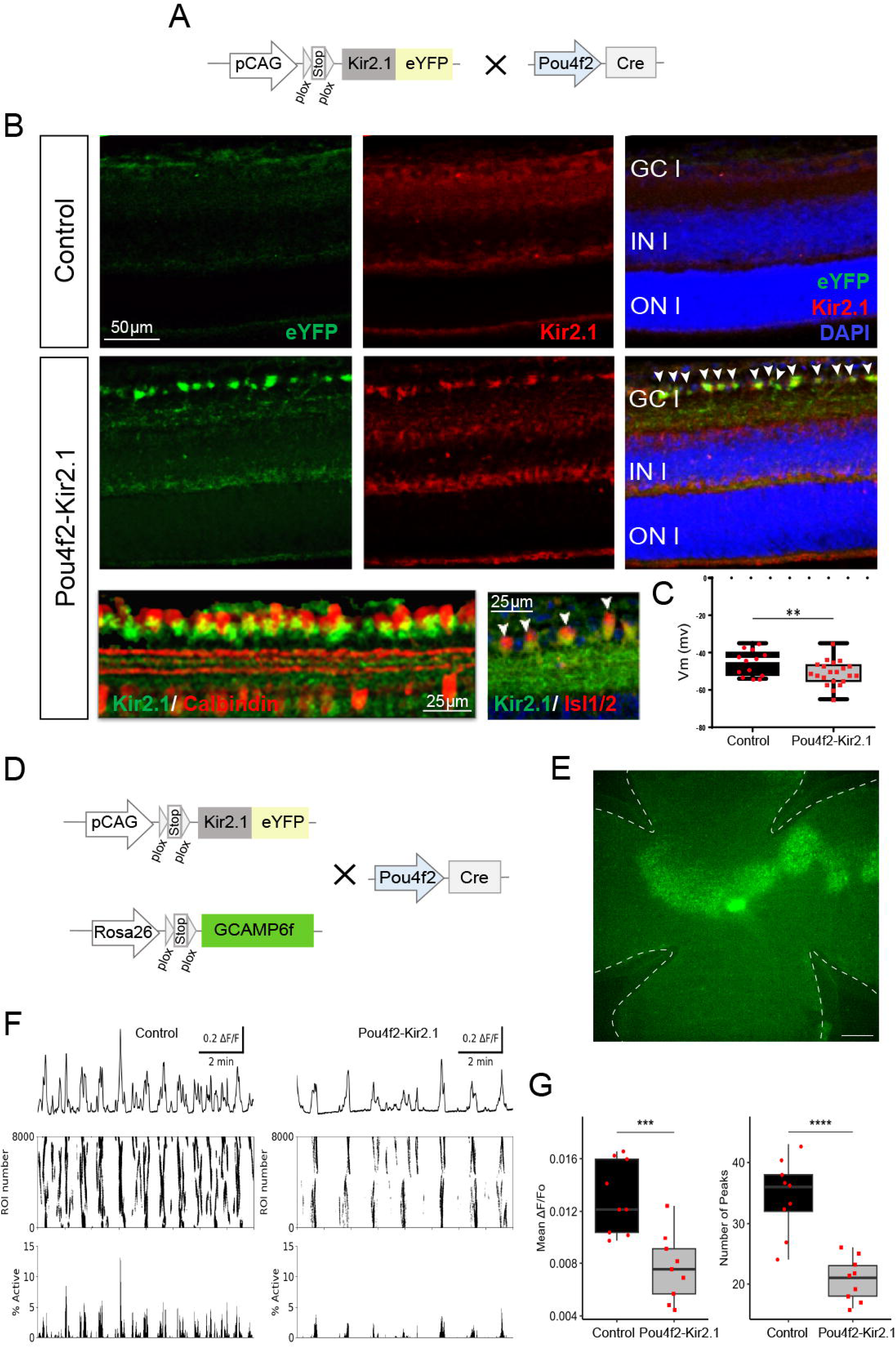
The frequency of retinal waves is lower in Pou4f2-Kir2.1 mice. **(A)** Generation of the Pou4f2-Kir2.1 mouse line. The Pou4f2-cre mice were breed with a conditional reporter line that contains a CAG-[Stop] cassette followed by the human Kir2.1 cDNA sequence fused to the eYFP coding sequence. **(B)** Immunostaining for Kir2.1 and YFP in retinal sections from adult Pou4f2-Kir2.1 and control littermate mice. Panels at the bottom show expression of Kir2.1 alone or combined with Calbindin in Pou4f2-Kir2.1 mouse retinas. **(C)** Averaged resting membrane potential in control cells and cells expressing Kir2.1. Dots represent TdTomato cells from retinas from three different mice. Whiskers extend to the min and max values (*p < 0.05, ** p<0.01, Student’s unpaired T-test). **(D)** Pou4f2-cre mice were breed with the Kir2.1 and the reporter Rosa26-GCaMP6f lines to obtain Pou4f2-Kir2.1;GCaMP6f mice. **(E)** Representative image of a wholemount flattened retina of a Pou4f2;GCaMP6f mouse. Retinal waves can be visualized on real time in the retinas of these mice. Scale bar: 300mm. **(F)** Top: Global average of ΔF/F traces representing calcium transients from all ROIs, in control and Pou4f2-Kir2.1 retinas. Middle: raster plot depicting calcium transients for all ROIs with ΔF/F signals greater than 50%. Bottom: percentage of ROIs active throughout the time course of the recording (10 minutes). **(G)** Box plots comparing the mean amplitude of the global fluorescence signal (left) and the number of waves detected as fluorescence peaks in 10-minute recordings (right), between control and Pou4f2-Kir2.1 retinas. Each data point represents a different animal. Statistical analysis revealed significant differences in both cases. For the fluorescence amplitude, the averages between the control and Pou4f2-Kir2.1 retinas were found to be statistically different. Similarly, for the number of peaks, the averages were also found to be statistically different. Whiskers extend to the min and max values (***p < 0.001, **** p<0.0001).

Next, to better visualize RGCs ectopically expressing Kir2.1 and patch clamp them, we breed Pou4f2-Kir2.1 mice with a reporter line (Rosa26^TdTm^). As expected, the resting membrane potential of RGCs recorded from postnatal Pou4f2-Kir2.1; Rosa26^TdTm^ mice (−50.7 ± 1.4 mV) was significantly lower than in neurons from control mice (Pou4f2-Rosa26^TdTm^; −45.5 ± 1.9 mV) (**Fig. 1C**). Then, to evaluate the impact of Kir2.1 ectopic expression in retinal waves, we crossed the Pou4f2-Kir2.1 mice with a cre dependent GCaMP6f calcium indicator reporter mouse line (Rosa26-stop^loxP^−GCaMP6f) which enables the visualization of retinal waves through fluorescence (**Fig. 1D**). We recorded semi-intact wholemount retinal preparations from P4 Pou4f2-Kir2.1; GCaMP6f mice and counterpart controls (Pou4f2-GCaMP6f) (**Fig. 1E**). Despite the ectopic expression of Kir2.1 in RGCs, we still observed correlated retinal waves but the frequency (peaks number) and amplitud of these waves were significantly lower compared to control retinas (**Fig. 1F-G, movies 1 and 2**).

Next, we analyzed the projection pattern of ipsi and contralateral RGC terminals at the dLGN and the SC by injecting cholera toxin subunit B conjugated with far-red (CTB-Alexa647) or green (CTB-Alexa488) fluorophores into each eye (**Fig. 2A**) of P9 Pou4f2-Kir2.1 and control (Kir2.1YFP^flxStop^) mice. Two days later, when axons are already refined but eyes still not open, anterograde axonal tracing was monitored in both iDISCO-clarified brains and coronal sections. As expected and in agreement with previous reports, segregation between ipsilateral and contralateral terminals was almost finished in the dLGN of control mice and a well-defined ipsilateral territory with no presence of contralateral innervation was clearly observed (Godement et al., 1984; Upton et al., 1999; Pham et al., 2001) (**Fig. 2B, 2D-D’ and 2G**). In contrast, contralateral terminals invaded the ipsilateral region in the dLGN of Pou4f2-Kir2.1 mice (**Fig. 2C, 2E-E’ and 2G**), resulting in an increased unsegregated area (**Fig. 2F**) and a concomitant reduction of the area innervated only by ipsilateral RGCs (**Fig. 2G**). Also, the area receiving ipsilateral innervation was expanded compared to the controls (**Fig. 2H**). The analysis of RGC axons at the SC also showed an aberrant refinement of ipsilateral RGC arborizations in Pou4f2-Kir2.1 mice compared to their control littermates (**Fig. S1**). Pou4f2 is also expressed in the SC and therefore, we cannot rule out that axon refinement defects in this nucleus result from activity alterations in the collicular cells. Despite this caveat in the SC, the lack of refinement in the thalamus confirms that a correct patterning of spontaneous retinal waves is essential for eye-specific refinement at the IFN and validate the Kir2.1YFP^flxStop^ mouse line as a model to investigate the impact of correlated spontaneous activity in other visual nuclei where Pou4f2 is not expressed.

**Figure 2.**
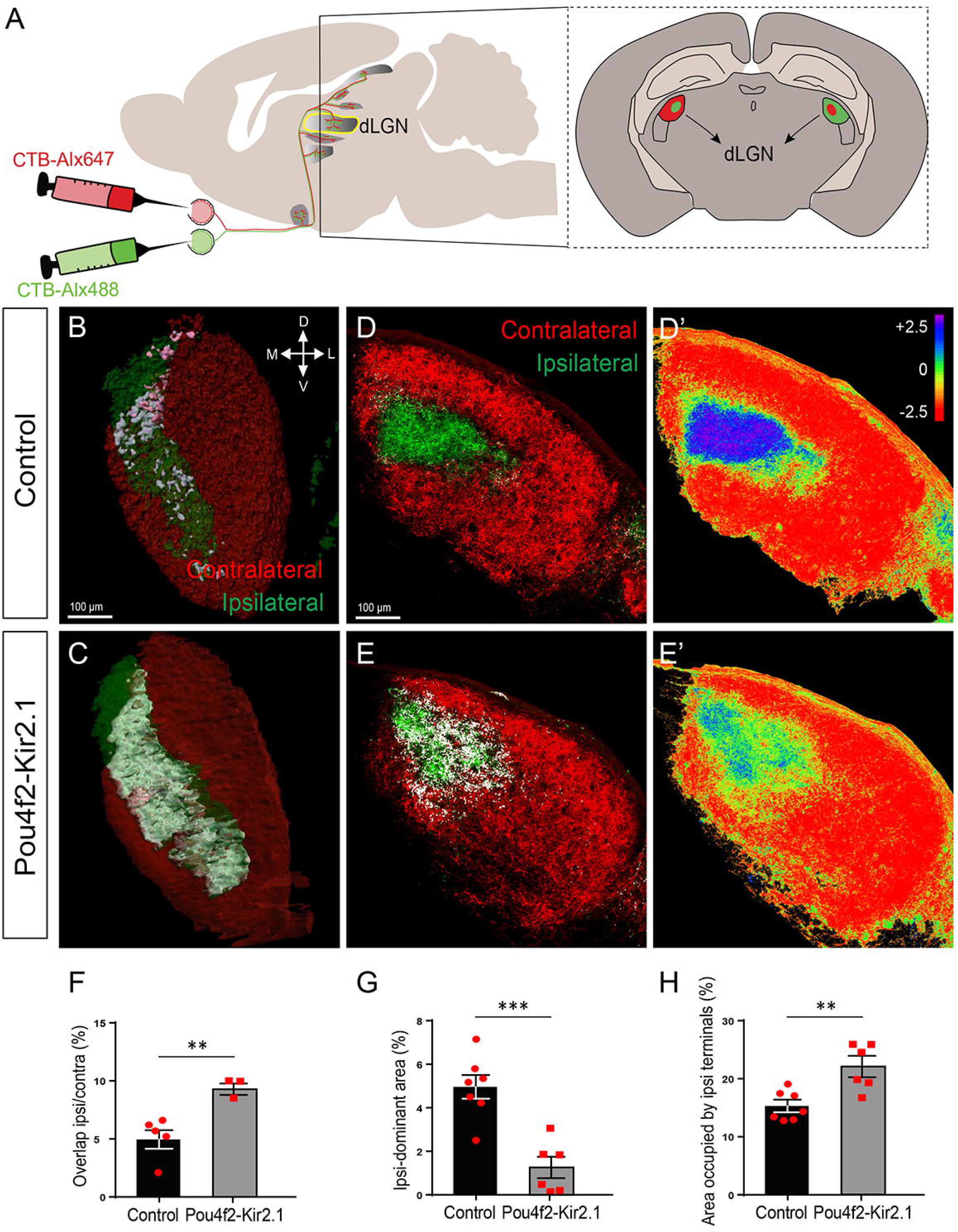
Pou4f2-Kir2.1 mice show eye specific segregation defects in the dLGN. **(A)** Retinal projections were labeled through intraocular injections of cholera toxin subunit B conjugated to different Alexa fluorophores (CTB-Alexa647 and CTB-Alexa488) for each eye. **(B, C)** 3D reconstructions and **(D-E)** coronal sections through the dLGN of P11 Pou4f2-Kir2.1 and control littermate mice injected with CTB-Alexa647 and CTB-Alexa488. Overlapping of ipsilateral and contralateral terminals is visualized in white. **(D’-E’)** R-distribution (log[ipsilateral/contralateral]) representing contralateral dominant area in red, unsegregated in green and ipsilateral dominant in blue. **(F)** Percentage of the area containing ipsilateral and contralateral terminals relative to the total area occupied by RGC terminals at the dLGN. **(G)** Percentage of the area with a majority of ipsilateral terminals relative to the total area occupied by RGC terminals (see the Materials and Method section). **(H)** Percentage of the area occupied by ipsilateral projections related to the total area occupied by RGC terminals. Each biological replicate (red dots) represents an animal. Error bars indicate ±SEM (** p<0.01, ***p < 0.001, Student’s unpaired T-test).

#### Transcriptional changes governing spontaneous-dependent refinement in the visual targets

Together with previous reports, our results indicate that retinal waves and the associated calcium influx into the cytoplasm influence the expression of genes that integrate the machinery required for axonal refinement and synapse formation (Bito et al., 1997; West et al., 2001). However, the molecular programs that act downstream of spontaneous activity to mediate RGC axon remodeling at the target nuclei have not been characterized. To identify molecular mechanisms underlying activity dependent-axonal remodeling, we took advantage of the Kir2.1 conditional mice and compared the transcriptional profile of RGCs from mice expressing Kir2.1 and control mice. For this, we crossed the Pou4f2-Kir2.1 with the reporter mouse line R26^Td^Tomato (Pou4f2-Kir2.1; R26^Td^Tm) and analyzed their RGCs compared to the control littermates (Pou4f2-; R26^Td^Tm) at two time points, P4, when most RGC axon arbors are remodeling at the visual nuclei and P8, when the majority of the terminals have already refined (**Fig. 3A**). Tomato labelled RGCs from Pou4f2-Kir2.1; R26^Td^Tm or control mice were purified by fluorescence activated cell sorting and their transcriptomes profile determined by high-throughput sequencing (RNA-Seq). Principal Component Analysis (PCA) confirmed differential grouping of samples from each genotype at both stages (**Fig. S2**).

**Figure 3.**
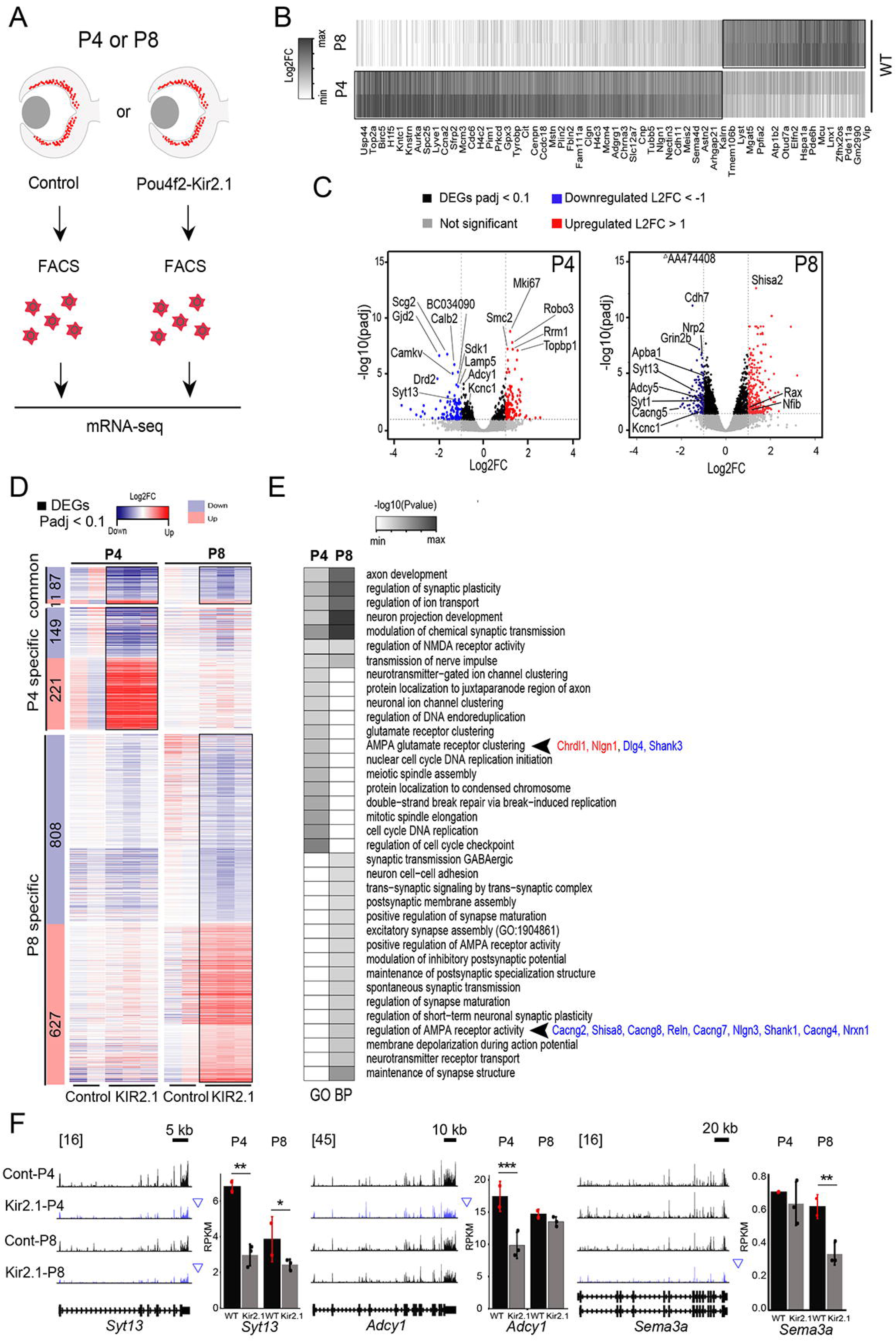
Identification of activity-dependent transcriptional programs underlying axon remodeling during axon refinement. **(A)** Scheme of the experimental approach. RGCs from P4 Pou4f2-Kir2.1 and control littermate mice were purified by FACS and their total RNA extracted to compare their transciptomic profiles. **(B)** Heatmaps of gene transcriptional profiles of RGCs from Pou4f2-Kir2.1 and control mice highlighting common and specific DEGs at P4 and P8. **(C)** Volcano plots showing the significance value distribution after DEGs analysis between Pou4f2-Kir2.1 and control mice at P4 and P8 (Padj < 0.1); |LFC|>1 = 217 genes. **(D)** Heatmaps of upregulated and downregulated DEGs in P4, P8 or common to both stages comparing Pou4f2-cre-Kir2.1 and control retinas. **(E)** Biological processes GO enrichment analysis of DEGs from genes with altered expression in P4, P8 or both in Pou4f2-cre-Kir2.1 mice compared to controls. P.adj < 0.1. **(F)** Gene profiles and graphs representing transcripts expression obtained in Pou4f2-Kir2.1 and control mice corresponding to three DEGs, *Adcy1*, Sema6a and *Syt13* at both stages.

The analysis of transcriptional profiles from P4 and P8 Pou4f2-Kir2.1; R26TdTm mice and their respective controls (**Fig. 3B**) revealed distinct deregulated transcriptional programs at each stage. The number of genes differentially expressed (DEG) between mutant and control retinas was lower at P4 (468) than at P8 (1533). At p4, 232 were upregulated and 236 downregulated while 895 were downregulated and 638 upregulated at P8 (**Fig. 3B, C and Table 1**). These differential temporal profiles likely reflect the switch from cholinergic to glutamatergic-dependent axonal remodelling known to occur during this time window. In a Gene Ontology (GO) analysis for Biological Processes, terms such as *Synapse Formation and Clustering of AMPA Receptors* (including genes as *Chrdl1, Nlgn1, Dlg4* and *Shank3* that encode proteins essential for synaptogenesis) were specific for P4. However, at P8 we retrieved terms related to more mature synapse stages such as *Synaptic Assembly and Maintenance*, *Regulation of NMDA and AMPA Receptor Activity*, or *Transmission of Spontaneous Activity*. Among the DEGs included in these terms we found members of the Type I transmembrane AMPA receptors (Cacng2/4/7/8), Shisa8, involved in the regulation of AMPA receptors and Nrxn1, that is known to play a role in the formation and maintenance of synaptic juctions via their calcium dependent interactions with Neuroligin family members (**Fig. 3D**).

Among the genes exclusively deregulated at P4 we found *Adcy1*, that encodes for Adenilate Cyclase 1, an enzyme involved in the refinement of visual axons at the dLGN and the SC (Nicol et al., 2006). In the set of DEGs specifically deregulated at P8 we found Sema3B, known to play an important function in neuroplasticity regulation right after eye opening (Carulli et al., 2021). Among the DEGs highly downregulated at both stages we found Syt13, that encodes for Synaptotagmin 13, a type I membrane protein involved in vesicular trafficking, exocytosis, and secretion (Fukuda and Mikoshiba, 2001; von Poser and Südhof, 2001) (**Fig. 3E**). Since Syt13 was deregulated at both stages, we aimed to manipulate the expression of this gene *in vivo* as a proof-of-concept to validate our screening. First, we analyzed the expression pattern of *Syt13 mRNA* by *in situ* hybridization in retinal sections from postnatal mice and confirmed that *Syt13* transcripts are highly and specifically expressed in P4 RGCs and less expressed after eye opening (**Fig. 4A**). Then, to investigate the function of Syt13 in axon refinement, we electroporated short-hairpin RNAs against Syt13 (*shRNA Syt13*) or control shRNA (*scramble shRNA*) together with plasmids encoding for the green fluorescence protein (GFP) into embryonic retinas (**Fig. 4B**). The analysis of targeted-RGC arborizations in sagittal sections of the SC before eye-opening (P10) showed very significant differences in the SC of *shRNA Syt13* compared to control (*shRNA scramble*) electroporated mice. Most labeled axons reached their corresponding topographical area at the SC in both control and *shRNA Syt13-expressing* electroporated mice. However, while control axons exhibited exuberant arborization in the target area, axons electroporated with shRNA Syt13 showed a significant decrease in arborization density (**Fig. 4C-D**). These results confirm a role for Syt13 in the refinement of RGC terminals at the SC and validate our unbiased screening designed to search for spontaneous activity-dependent molecules involved in axonal remodeling at the visual targets.

**Figure 4.**
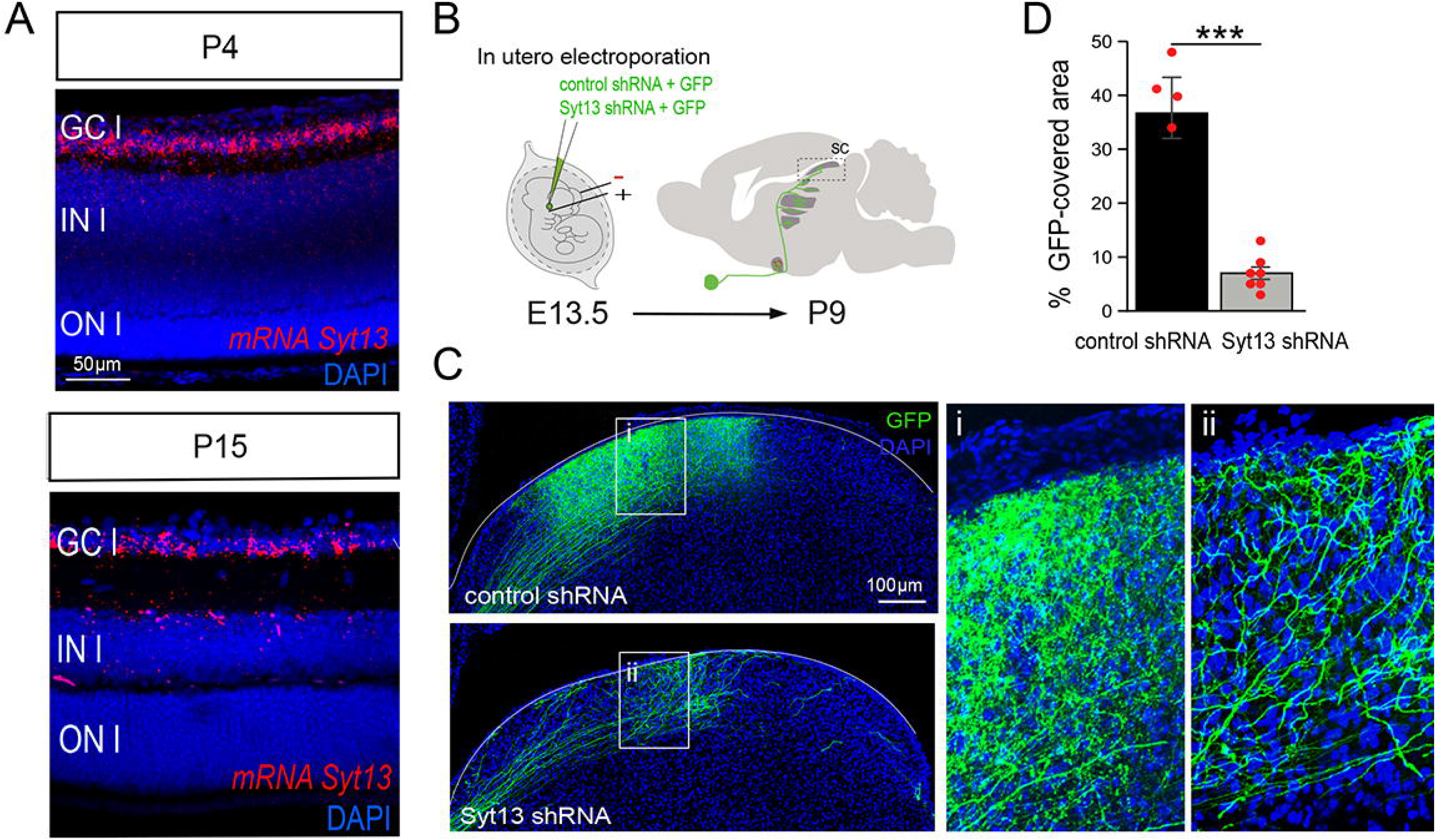
Syt13 is essential for axon refinement at the SC. **(A)** Representative image of a retinal section from P4 and P15 wildtype mice showing *in situ* hybridization for Syt13. GCl, Ganglion cell layer, INl, Inner layer, ONl, outer layer. **(B)** Schematic representation of the experimental approach. **(C)** Representative examples of sagittal sections through the Superior Colliculus (SC) of P11 mice electroporated at E13 with plasmids encoding for GFP and control shRNA or Syt13shRNA and contrastained with DAPI. Right panels (i, ii) show high magnification images from the squared areas. **(D)** Fluorescence intensity (FI) from axons of RGCs targeted at E13 by electroporation measured in sagittal sections at the SC level and normalized to the total area occupied by the SC in each slide. Red dots represent percentage of area covered by electroporated axons (see the Material and Methods section) obtained by averaging results from two central sagittal sections of one SC/animal (Student’s unpaired T-test, ***p-value < 0.001). Results show means ± SEM.

#### Altered retinal waves affect axon terminals refinement at the NIFN

Correlated retinal activity is known to encode spatial information among neighboring RGCs. Since NIFN are not as functionally-related to spatial information and sight as IFN, we wondered to what extent, if at all, the alteration of spontaneous retinal waves affects the refinement of visual terminals in these nuclei. To address this, we analyzed axonal terminals at two NIFN, the olivary pretectal nucleus (OPN) and the suprachiasmatic nucleus (SCN).

The OPN is the largest of the seven nuclei that integrate the pretectal nuclei located in a region anterior to the SC. It is involved in mediating behavioral responses to acute changes in ambient light and the optokinetic reflex (Gamlin, 2006). We analyzed serial coronal sections from Pou4f2-Kir2.1 and control mice previously injected with CTB-Alexa647 or CTB-Alexa488 into each eye (**Fig. 5A**). A larger number of ipsi and contralateral terminals overlapped in space in the OPN of Pou4f2-Kir2.1 mice compared to control littermates (**Fig. 5B-F**). These results demonstrate segregation defects in this NIFN as a consequence of disturbed retinal spontaneous activity.

**Figure 5.**
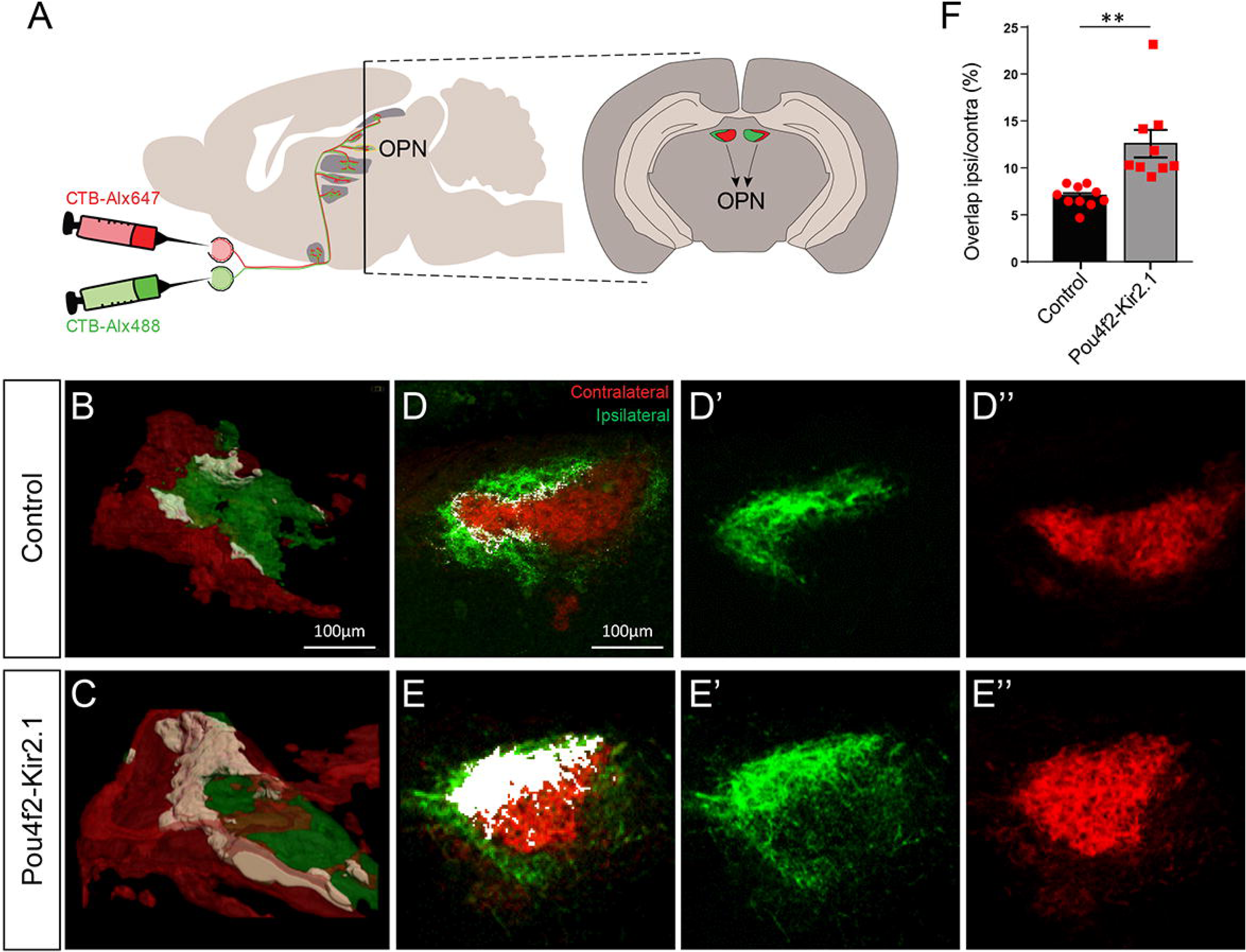
Proper frequency of retinal waves is essential for eye-specirid segregation in the OPN. **(A)** Retinal projections were labeled through intraocular injections of cholera toxin subunit B fused to different Alexa fluorophores (CTB-Alexa647 and CTB-Alexa488) for each eye. **(B-C)** 3D reconstructions and **(D-E)** coronal sections through the OPN of P11 Pou4f2-Kir2.1 and control littermate mice injected with CTB-Alexa647 and CTB-Alexa488. Overlapping of ipsilateral and contralateral terminals is visualized in white. **(F)** Percentage of the area containing ipsilateral and contralateral terminals (white) relative to the total area occupied by RGC terminals. Each biological replicate (red dots) represents an animal. Error bars indicate ±SEM (**p<0.001, Student’s unpaired T-test).

The SCN is located in the diencephalon just behind the optic chiasm. This NIFN nucleus, considered the circadian pacemaker, is responsible for controlling circadian rhythm and it is innervated only by intrinsically photosensitive RGCs that express melanopsin (OPN4), distribute all over the retina (Sekaran et al., 2003; Prigge et al., 2016) and can be activated by the retinal waves (Renna et al., 2011). Most Pou4f2-RGCs do not project to the SCN (Chen et al., 2011). Thus, to investigate the function of retinal spontaneous activity in this nuclei, we used a second cre-line (Slc6a4/Et33-Cre) that drives gene expression to a subset of RGCs that mainly project to the ipsilateral IFN (García-Frigola and Herrera, 2010) but also innervate the SCN (Su et al., 2021).

To visualize Slc6a4-positive RGCs and their terminals in the SCN nuclei, we bred Slc6a4/Et33-Cre mice with a tdTomato reporter line. While intrinsically photosensitive retinal ganglion cells make up approximately 2% of the total RGC population (Berson et al., 2002; Hattar et al., 2002; Do and Yau, 2010; McNeill et al., 2011; Schmidt et al., 2011), we made a noteworthy observation of a twofold enrichment of OPN4-positive cells (~4%) within the Slc6a4-RGCs population (**Fig. 6A**). Then, we analyzed the distance between axon terminals in the SCN coming from the ipsilateral or contralateral eye. Far from being segregated, we noticed that after injecting CTB-Alexa488 and CTB-Alexa647 into the eyes of Slc6a4-TdTm mice, about 90% of the Slc6a4-terminals coming from one eye (tdTm/CTB-Alexa 488) were located only 5μm apart from axons arising in the opposite eye (CTB-Alexa647) (**Fig. 6D**). This result confirms previous studies showing that melanopsin RGC terminals do not undergo eye-specific segregation in the SCN and different subtypes of SCN cells indeed receive input from both retinas (Fernandez et al., 2016).

**Figure 6.**
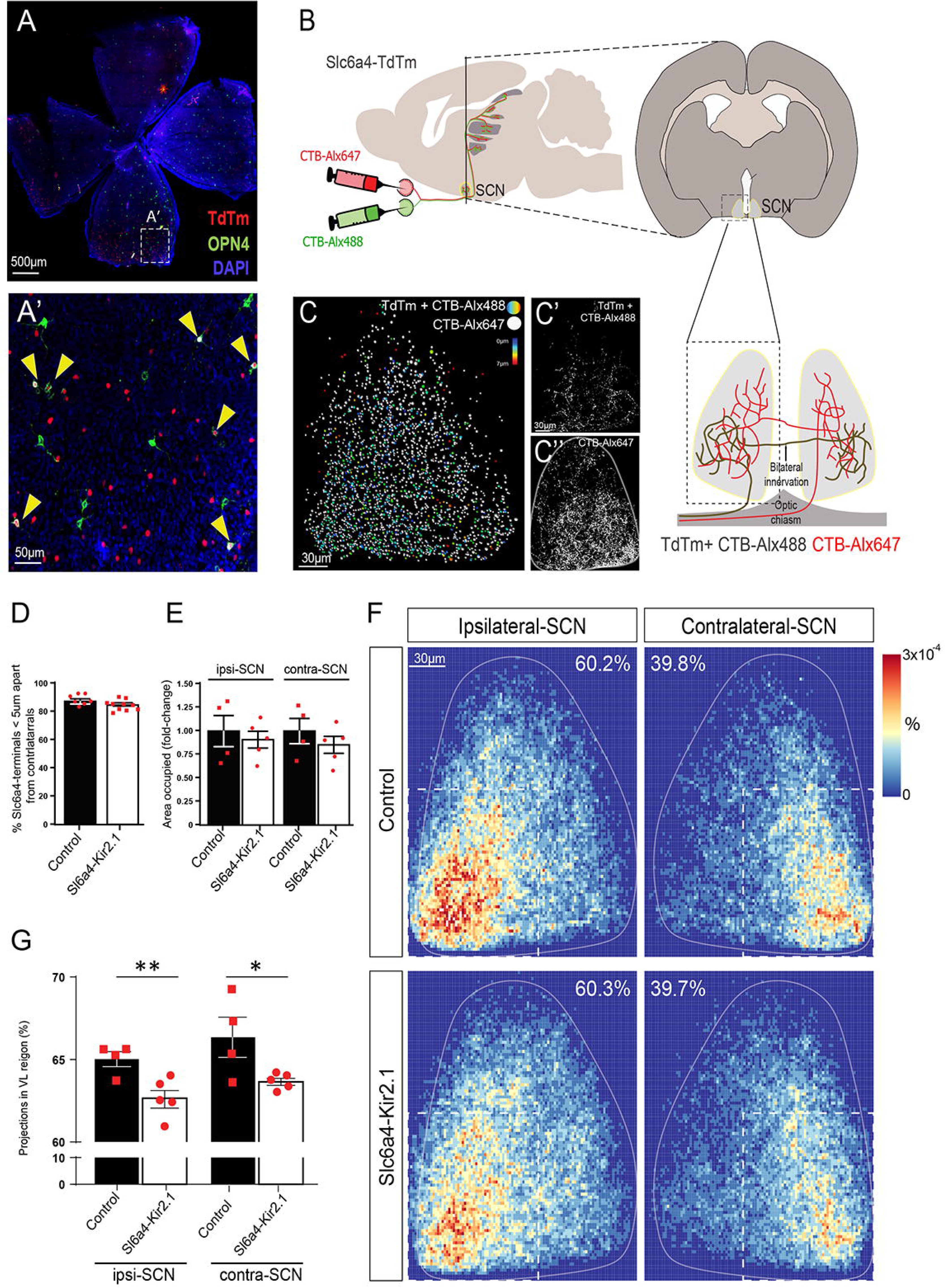
Optimal frequency of retinal waves is needed for the refinement of visual axons in the SCN. **(A-A’)** Whole-mount retina from P11 Slc6a4-TdTm mice stained with αOPN4 and αtdTomato antibodies. Yellow arrowheads point to double stained cells. **(B)** Injections of CTB fused to Alexa-488 or Alexa-647 in P11 Slc6a4-TdTm mice and analysis of coronal sections at the level of the SCN (yellow) two days later. Retinal projections from each eye bilaterally innervate the SCN. **(C)** Representation of retinal terminals from control mice displaying ipsilateral terminals colored according to their distance to the nearest contralateral neighbor. Contralateral terminals are showed in gray. **(C’-C’’)** Single Z from the stack used in C showing a representative binary mask for ipsilateral (C’) or contralateral (C’’) terminals. **(D)** Percentage of ipsilateral terminals closer than 5µm to a contralateral. **(E)** Area occupied by TdTm+ retinal terminals comparing control and Slc6a4-Kir2.1 mice (fold-change) in both sides of the SCN. **(F)** Density maps displaying projections from the ipsilateral RGCs to the SCN in Slc6a4-Kir2.1 (N=5) and control (N=4) mice, obtained by averaging all the replicates per condition. Percentages in the top center indicate the ratio of ipsilateral RGC innervation to each side of the SCN. **(G)** Ratio of projections innervating the ventrolateral quadrant of the SCN for each replica on both sides of the SCN. Red dots represent biological replicates. Error bars indicate ±SEM (*p < 0.05, ** p<0.01, Student’s unpaired T-test).

To then investigate the influence of spontaneous waves in the wiring of visual input at the SCN, we injected Alexas into the eyes of Slc6a4-Kir2.1 and control littermate mice and first confirmed that, compared to the controls the resting membrane potential of RGCs recorded from postnatal Slc6a4-Kir2.1; Rosa26^TdTm^ mice (−55.1±1.5 mV) was significantly lower than in neurons from control mice (Slc6a4-Kir2.1-Rosa26^TdTm^; −46.4 ± 2.6 mV) and that the RGC axon terminals of Slc6a4-Kir2.1; Rosa26^TdTm^ mice do not refine properly at the LGN and the SC (**Fig. S3**) indicating that retinal activity is also perturbed in these mice. Next, we analyzed Slc6a4-terminals at the SNC by analyzing TdTm/CTB-Alexa488 at the SCN of Slc6a4-Kir2.1 and control mice. TdTm/CTB-Alexa488 arborizations covered similar areas in Slc6a4-Ki2.1 and control mice (**Fig. 6E**). However, the analysis of the density maps demonstrated that axonal arborizations were significantly reduced in the Slc6a4-Kir2.1 mice compared to the controls in the ventrolateral area of this nucleus (**Fig. 6F, G**). These results strongly suggested that, despite eye-specific segregation and topographic mapping are not evident features of the SCN, spontaneous retinal activity before eye-opening influences the innervation of RGCs terminals in this NIFN.

## DISCUSSION

Spontaneous neural activity is essential in a multitude of processes in the developing nervous system. Spontaneous calcium transients have been detected in neocortex and thalamus in prenatal stages and may modulate neurogenesis wiring and neurotransmitter identity (Borodinsky et al., 2004; Corlew et al., 2004; Bonetti and Surace, 2010; Antón-Bolaños et al., 2019). In the visual system, correlated spontaneous retinal activity is known to be essential for the fine-tune of retinotopic maps and eye-specific segregation in the IFN, but it was unclear to what extent the correct connectivity of RGC axons at NIFN also depend on retinal waves. In this study, by generating several mouse lines with disturbed retinal waves before eye opening and analyzing the refinement of RGCs axon terminals in the SCN and the OPN, we found that an optimal frequency of retinal waves in pre-eye opening stages is essential for the establishment of proper connectivity also in NIFN. In addition, we retrieved the transcriptional programs triggered by retinal activity essential for the proper assembly of the visual circuit.

### Ectopic expression of Kir2.1 in RGCs alters the frequency of retinal waves

The rectifying potassium channel Kir2.1 is a known suppressor of intrinsic excitability, which has a crucial role in establishing the connectivity of several circuits. Overexpression of Kir2.1 has been used to electrically silence neurons in both *in vitro* and *in vivo* experiments (Burrone et al., 2002; Benjumeda et al., 2013). Moreover, ectopic expression of Kir2.1 in transgenic mice has been also reported to block neural activity in the olfactory system (Yu et al., 2004) and may lead multiple different effects in vivo, depending on the developmental stage of the network (Burrone et al., 2002; Antón-Bolaños et al., 2019). We observed that recombinant RGCs in the Kir2.1 mice exhibit a more negative resting potential than RGCs in the control mice. This translated into a lower frequency of retinal waves, which is sufficient to alter the refinement of retinal projections at the different visual targets.

### Transcriptional programs involved in activity-dependent axonal remodeling

Pou4f2-Kir2.1 mice show significant defects in fine-scale/local refinement of axonal arborizations at the different visual nuclei, confirming that the mechanisms involved in the remodeling of visual axons are influenced by retinal waves. By comparing the transcriptomic profile of Pou4f2-Kir2.1 and control retinas at P4 and P8 we identified genes (DEGs) potentially regulated by retinal activity during postnatal stages before eye-opening. Interestingly, these DEGs seem to reflect the transcriptional changes associated to the transition of RGCs switching from a scenario dominated by axon branching dynamics to an assembled circuit carrying out synaptic maturation. Thus, among the P4 DEG specific genes we found *Adcy1*, which was already described as important for retinal axon refinement, and *Nlgn1,* which encodes for a protein important for the recruitment and clustering of other proteins during synaptogenesis. At the later stage (P8), when connections are more mature, we found Nrxn1 deregulated in the Kir2.1 mutant retinas. Neurexins and Neuroligins form complexes at the synapses during synaptogenesis and our results suggest that the expression of these genes that encode for proteins essential for synapse formation depend, at least partially, on proper patterns of retinal waves.

Interestingly, we also found a number of DEGs related to cell cycle. It has been recently reported that beyond its role in DNA damage response, the serin/threonine kinase ATR plays a function in regulating neuronal activity by modulating presynaptic firing. ATR interacts with synaptotagmins in synaptosomes and we found Synaptotagmin13, one of the less studied members of the synaptotagmin family (Wolfes and Dean, 2020), altered in the Kir2.1 mice. This could explain, at least in part, the *double-strand bread repair visa break-induced replication* GO terms that we retrieved in our analysis. It is important to mention that unlike other members of the Syt family, Syt13 does not sense calcium (Bhalla et al., 2008). However, we observed that Syt13 downregulation leads to altered axonal arborizations at the SC. These results revealed a novel role for this poorly known member of the Syt family but further experiments are needed to precisely determine the function of this, and other candidates retrieved in our screening, in the remodeling of RGC axon terminals.

### Particularities of retinal activity-dependent refinement in NIFN

A large number of studies have demonstrated that ipsilateral and contralateral axon terminals initially overlap in the dLGN and are later expelled from the territory that corresponds to the other eye, competing with the contrary retinal input in terms of activity correlation. We observed eye-specific segregation defects in the IFN of both Pou4f2-Kir2.1 and Slc6a4-Kir2.1 mice. In this study we show that, as in the IFN, when activity is altered, cRGCs terminals occupy the ipsilateral territory in a NIFN as the OPN. Thus, despite the different anatomies, features and functions of the OPN compared to IFN, RGC terminals appear to follow similar rules to target and refine visual nuclei in terms of eye-specific activity competition in nuclei that receive a number of retinal terminal sufficient to compite with axons from the other eye. In the SCN the scenario is different. Eye-specific segregation does not take place in this nucleus. RGC axons project bilaterally (Fernandez et al., 2016) and arborizations coming from each eye are likely not abundant enough to compete with those coming from the other eye. The SCN was thought to be just a regulator of the circadian clock. However, retinal input to the SCN encodes for a much more complex visual information than that necessary to just perform this function (Mouland et al., 2017; Stinchcombe et al., 2017; Yu et al., 2017), suggesting a potential role for this nuclei in spatial vision as well. Our experiments indicating that an optimal frequency in the number of retinal waves is required for the proper connectivity of this nucleus support this hypothesis. Future experiments in conditional mice with altered retinal activity in all melanopsin-RGCs should better define the role of spontaneous activity in the refinement of retinal input to the SCN as well as a potential impact in mice behavior.

## MATERIALS AND METHODS

### Mice

The Kir2.1eYFP^flx-Stop^ line was generated in our laboratory by cloning the human Kir2.1 cDNA (Benjumeda et al., 2013) in a pEYFP-N1 plasmid adding the eYFP reporter protein at the C-terminal of Kir2.1. This fusion protein was subcloned in the PCAG-SE plasmid downstream of the CAG promoter and a LoxP-flanked STOP cassette. The function and plasma membrane localization of the chimeric channel were confirmed by electrophysiology. The linearized and purified construct was injected in mice oocytes (CEBATEG, Barcelona). Twelve funders were obtained and crossed with Pou4f2-cre mice (Pou4f2^tm1(cre)Bnt^/J; RRID:IMSR_JAX:030357) to select the funder with the highest Kir2.1YFP expression in RGCs. Slc6a4-cre line belongs to the GENSAT Project at Rockefeller University (B6.FVB(Cg)-Tg(Slc6a4-cre)ET33Gsat/Mmucd; RRID:MMRRC_031028-UCD). Rosa26Stop-tdTm (B6.Cg-Gt(ROSA)26Sor^tm14(CAG-tdTomato)Hze^/J; RRID:IMSR_JAX:030357) and GCaMP6f mice (B6;129S-Gt(ROSA)26Sor^tm95.1(CAG-GCaMP6f)Hze^/J; RRID:IMSR_JAX:024105) were acquired from Jackson Laboratories. Animals were housed in a timed-pregnancy breeding colony at the Instituto de Neurociencias de Alicante, Spain. Conditions and procedures were approved by the IN-Animal Care and Use Committee and met European (2013/63/UE) and Spanish regulations (RD 53/2013).

### Electrophysiology

Retinas extracted from P4 or P11 (Pou4f2-Kir2.1;R26^Td^Tm) and control (Pou4f2; R26^Td^Tm) mice were placed in artificial cerebrospinal fluid solution (ACSF) at 4 ⁰C, containing (in mM): 126 mM NaCl, 26 mM NaHCO_3_, 10 mM Glucose, 5 mM MgCl_2_, 3 mM KCl, 1.25 mM NaH_2_PO_4_, 1 mM CaCl_2_. pH of the ACSF solution was tampered using carbogen (5% CO2 and 95% oxygen) and the resulting osmolarity was ~310 mOsmol/Kg. Further, retinas were subjected to enzymatic digestion with collagenase/dispase diluted in ACSF (1:3) for 15-20 min at 35°C with carbogen. Collagenase/dispase mix was prepared as follow: 155 mM NaCl, 1.5 K_2_PO_4_, 10 mM HEPES, 5 mM glucose), collagenase type XI (Sigma, D9542) 900 uni/ml, dispase (BDBiosciences, 10103578.001) 5.5 uni/ml. Recordings were performed using an Olympus BX50WI up-right confocal microscope and constant perfusion with ACSF solution at ~34 °C. Temperature was controlled by a feedback Peltier device CL-100 (Warner Instruments, Hamdem, CT, USA). Neurons were visualized through a 40x water immersion objective. To identify tdTm labeling, the field was expose to a 554 nm light with a Polychrome V monochromator (Thermo Scientific, USA) and the fluorescence emission captured on a cooled CCDge camera (sensicam, PCO AG, Kelheim, Germany). Neurons were current-clamped and once in the whole-configuration, resting membrane potential was measured. Recordings were performed using a Multiclamp 700B amplifier, pCLAMP 10 software and a Digidata 1440A and pCLAMP 10 software (Molecular Devices, Sunnyvale, USA). Borosilicate electrodes were pulled using a puller P-97 (Sutter Instrument, CA, USA) with resistance ranging between 4 to 8 MΩ and filled with an intracellular solution containing: 115 mM K-gluconate, 25 mM KCl, 9 mM NaCl, 10 mM HEPES, 0.2 mM EGTA, 1 mM MgCl_2_, 3 mM K_2_-ATP and 1 mM Na-GTP adjusted to pH 7.2 with KOH (280–290 mOsmol/kg). Data were sampled at a frequency of 20 KHz and low-passed filtered at 10 KHz. Statistical analyses were performed with SigmaStat 4.0 software (Systat Software) and Origin Pro8 software (OriginLab Corporation) was used for graphing.

### Retinal waves recordings and analysis

Wholemount retinas from P4 Pou4f2-Kir2.1;GCaMP6f and control (Pou4f2-GCaMP6f) mice were dissected in oxygenated (5% CO2 and 95% oxygen) artificial cerebrospinal fluid solution (ACSF) at room temperature, containing: 119 mM NaCl, 5 mM KCl, 1.3 mM MgSO_4_ 7H_2_O, 1 mM NaH_2_PO_4_, 26 mM NaHCO_3_, 11 mM Glucose, 2.4 mM CaCl_2_. Retinas were mounted over a Whatman filter paper (Whatman, WHA150446) with the ganglion cell layer side up and transferred turned upside down to a cell culture insert (Millipore, PICM0RG50) in a MatTek glass bottom dish (MatTek, P35G-1.5-14-C) with 1mL of ACSF.

Then, the dishes with the retinas were placed in a warmed (32°C) and gassed recording chamber of an inverted Leica Thunder Imager microscope. After 20 minutes in the chamber, time-lapse spontaneous calcium recordings were obtained with a 5x dry objective (5x/0.12 dry WD = 14mm) after exciting the retinas at 475 nm. Images were acquired with a DFC 9000 GTC sCMOS camera for 10 minutes using a time interval of 300ms, an exposure time of 200ms, a LED intensity of 50% and 4×4 binning.

For the analysis, retina.lif videos were read with Fiji (Schindelin et al., 2012). The mean value of the fluorescence as a function of time was obtained with the Fiji function Plot Z-axis Profile, and saved as .csv files for further analysis with Python. The fluorescence peaks corresponding to the presence of calcium waves were detected with the SciPy function scipy.signal.find_peaks (Virtanen et al., 2020).

For the analysis of regions of interest (ROIs), 3×3 pixel areas (corresponding to 15.6×15.6 μm) were drawn onto the retina image. These ROIs were separated by 2 pixels (corresponding to 10.4 μm). The average fluorescence intensity value was computed for all frames and the resulting signal was detrended and normalized. Additionally, a global intensity signal was obtained by averaging the signals from all ROIs.

### Anterograde labeling of retinal projections

Postnatal mice were anaesthetized in ice or with isofluorane depending on the age. Whole-eye anterograde labelling was performed using a *Nanoject II Auto-Nanoliter Injector (Drummond, 3-000-204)* injecting twice on opposite points with 0.5µl of CTB-subunit conjugated with Alexa Fluor 488 or 647 *(Invitrogen, C34775/C22843/C34778) as* described in (Rebsam et al., 2009). Two days later mice were perfused transcardially with 4% paraformaldehyde (PFA) in 0.1 M phosphate-buffered saline (PBS). Brains were incubated overnight in the PFA solution at 4°C and stored in PBS. Fixed tissue was embedded in 3% agarose diluted in PBS and sectioned in a vibratome. Sections were incubated in blocking solution (0.2% gelatin, 5% FBS, 0.005-0.01% Tx) at RT for 1h and mounted to be analyzed.

### Inmunofluorescence and in situ hybridization

Sectioned PFA fixed tissue was incubated in blocking solution (0.2% gelatin, 5% FBS, 0.005-0.01% Tx) at RT for 1h, followed by incubation O/N with the primary antibodies. The next primary antibodies were used at the specified concentrations: rabbit anti-Dsred Pab (Takara Bio Cat# 632496, RRID:AB_10013483) [1:500], rabbit polyclonal Kir2.1 (Abcam Cat# ab65796, RRID:AB_1140953) [1:500], chicken anti-GFP (Aves Labs Cat# GFP-1020, RRID:AB_10000240) [1:1000], rabbit polyclonal anti-melanopsin (Advanced Targeting Systems Cat# AB-N38, RRID:AB_1608077) [1:2500]. After three washes in PBS, samples were incubated with the secondary antibodies diluted 1:1000 in blocking solution at room temperature for 2 h. For whole-mount retinas, the protocol was the same used for sections but Triton concentration was increased to 1% and samples were fixated in methanol. This process includes washes with 25%-50%-80%-100% methanol/H2O for 20’ at RT, a bleaching in 3% H2O2 in methanol for 1h at RT, two washes in methanol, rehydration in 80%-50%-25% methanol/PBS 20’ each at RT and blocking in 5% BSA PBST (3% Tween) for 1 hour at RT.

In situ hybridization for *mRNA Syt13* was performed according to reported methods (Murcia-Belmonte et al 2019). The mouse probe (688bp) was amplified from mouse cDNA using the following primers: Forward 5’-GAAACACCAGGCTCAGAAGC-3’; Reverse 5’-ACTGAGGGTACCTGGCACAT-3’.

### Brain clarification

Clarification was performed following the iDISCO protocol (Renier et al., 2014). Briefly, samples were dehydrated in methanol/H2O series (20%-40%-60%-80%-100%-100%) 1h at RT. Then samples were incubated 3h, with shaking, in 66% DCM/methanol at RT, and washed in 100% DCM to wash the methanol. Finally, brains were incubated and storaged in a glass tube totally filled with DiBenzylEther at RT. Brains showing total filling of contralateral projections labelled with CTB-546 and ipsilateral with CTB-647 were selected. For 3D quantifications, dLGN stacks were previously transformed to IsoData binary masks in ImageJ (IsoData algorithm) to standardize rendering in Imaris (Bitplane).

### Image acquisition and analysis

Images from tissue sections were captured using Olympus FV1000 confocal IX81 microscope and FV10-ASW software (Olympus). Acquisitions from clarified brains were made using a Light Sheet microscope (LaVision Ultramicroscope II) and LaVision BioTec Inspector Pro (LaVision). 3D rendering and processing were carried out in Imaris 9.1.2 (Bitplane). Images were processed in the ImageJ distribution Fiji (Schindelin et al., 2012) in order to denoise, enhance, threshold, co-localize or measure areas and distances, depending on the type of quantification. In all cases, background fluorescence was subtracted from sections using a rolling ball filter and gray scale was renormalized so that the range of gray-scale values was from 0 to 256. Data obtained from the image analysis were loaded into R 3.6.0 (R Core Team 2019) to perform mathematical calculations and generate some of the graphs. Statistical analysis and the rest of graphs were carried out using GraphPad Prism version 6.00 (GraphPad Software). Figures and graphs were edited with Adobe Illustrator CS6 (Adobe Systems Inc.). Error bars indicate ± SEM. (**p < 0.01, ***p < 0.001, Student’s unpaired t test).

### Eye-specific segregation analysis

Binocular co-localization was computed overlapping the “IsoData” binary mask of the Z-stack for each channel. We selected the two (OPN, SCN) or three (dLGN) most central 70-micron sections from the nucleus. The result was normalized by the total area of the nucleus. R-distribution was carried out as described in Torborg, et al. (Torborg and Feller, 2004) computing the logarithm of the intensity ratio (*R* = *log*_10_ (*F_I_*/*F_C_*)). The intensity threshold was set at 5 and empty pixels in both channels were discarded. The remaining zero-value pixels were replaced by 0.01 to allow the calculation of the logarithm. Range [-0.5, +0.5] was considered non-segregated area and [+1.75, +2.5] as ipsidominant area. The ipsilateral territory was obtained by dividing the pixels with ipsilateral signal to the total number of pixels of the dLGN.

### Suprachiasmatic nucleus density and proximity analysis

All 60-micron coronal sections of the SCN were analyzed, being aggregated by sample and averaged by group. Since retinal projections to the SCN project bilaterally, on each side of the nucleus we can find terminals labeled with the same CTB-Alx conjugate coming from opposite retinas. Therefore, TdTm pixels co-localizing with both Alexa fluorophores were discarded because of the inability to assign them to a specific retinal input. For the ipsilateral/contralateral proximity analysis, regions with high CTB-Alexa accumulation were selected as an approximation to synaptic boutons. Ipsilateral boutons are those labeled by the corresponding CTB-Alexa conjugate and tdTm. Contralateral boutons are those positive for the other CTB-Alexa conjugate and lacking tdTm signal. In the computation of the density maps, terminals were measured by summing the binary mask for each Z-plane along all sections of the SCN, combining and averaging the results for each source retina based on the corresponding CTB-Alexa conjugate (Alexa Fluor 488 or 647). Samples were normalized for nucleus size and signal intensity prior aggregating by group to generate a density map for each SCN side per condition.

### RNA-seq and analysis

Retinas were dissected out from P4 Pou4f2-cre; R26^Td^Tomato (control RGCs) and Pou4f2-Kir2.1; R26^Td^Tomato (Kir2.1 RGCs) mice. Retinas for each genotype were isolated and enzymatically dissociated in a mixture of collagenase/Trypsin and 1% BSA for 20 minutes at 37.C followed by mechanical dissociation. A 40 μM cell strainer was used for aggregates removal after dissociation and TdTm-expressing RGCs were isolated by fluorescent-activated cell sorting (BD FACS Aria™ III. BD Bioscience). Total RNA extraction was performed with Arcturus Picopure RNA isolation Kit (ThermoFisher Scientific). Three independent samples containing total RNA from 9-11 pooled retinas from at least three different litters/condition were sequenced according to the manufacturer instructions in a HiSeq Sequencing v4 Chemistry (Illumina, Inc). Briefly, RNAseq reads were mapped to the mouse genome (Mus_musculus.GRCm.38.83) using STAR (v2.5.0c) (Dobin et al., 2013). Quality control of the raw data was performed with FastQC (http://www.bioinformatics.babraham.ac.uk/projects/fastqc/). Library sizes were between 17 and 23 million single reads. Samtools (v1.3.1) was used to handle BAM files (Li et al., 2009). To retrieve differentially expressed genes (DEG), mapped reads were counted with HTSeq v0.6.1 (Anders et al., 2015) with the following parameters: -s reverse --i gene_id and with a gtf file reference for GRCm38.83. Samples were adjusted for library size and normalized with the variance stabilizing transformation (vst) in the R statistical environment using DESeq2 (v1.10.0) pipeline (Love et al., 2014). When performing differential expression analysis between groups, we applied the embedded IndependentFiltering procedure to exclude genes that were not expressed at appreciable levels in most of the samples. A heatmap of this distance matrix gives us an overview over similarities and dissimilarities between samples with a clustering based on the distances between the rows/columns of the distance matrix (FigS4). The PCA is generates using the most variable genes detected in the entire dataset. DESeq2 default values are set to use only the 500 most variable genes. This number is often applied when transcription of protein-coding genes is analysed (FigS4). Gene expression values were represented using the normalization techniques provided by each algorithm: *Reads per Kilobase of Mapped reads* (RPKM), *Relative Log Expression* (rlog from *DESeq2*). Analysis and preprocessing of data were performed with custom scripts using R (https://cran.r-project.org/) (v3.4.3) statistical computing and graphics, and bioconductor v3.2 (BiocInstaller 1.20.3) (Huber et al., 2015). Genes were considered differentially expressed at Benjamini–Hochberg (BH) adjusted, Padj < 0.1 and abs|Log2FC| > 1. Significantly upregulated and downregulated genes were visualized with IGV (v2.3.72) (Thorvaldsdóttir et al., 2013). We performed an Enrichment Analysis for Gene Ontology using the platform Panther (Mi et al., 2019) Fisher’s exact test and the Padj correction were used, obtaining the top terms using the filters by ratio enrichment > 2, number of GO family group genes between 3 and 2000, number of enrichment genes >3.

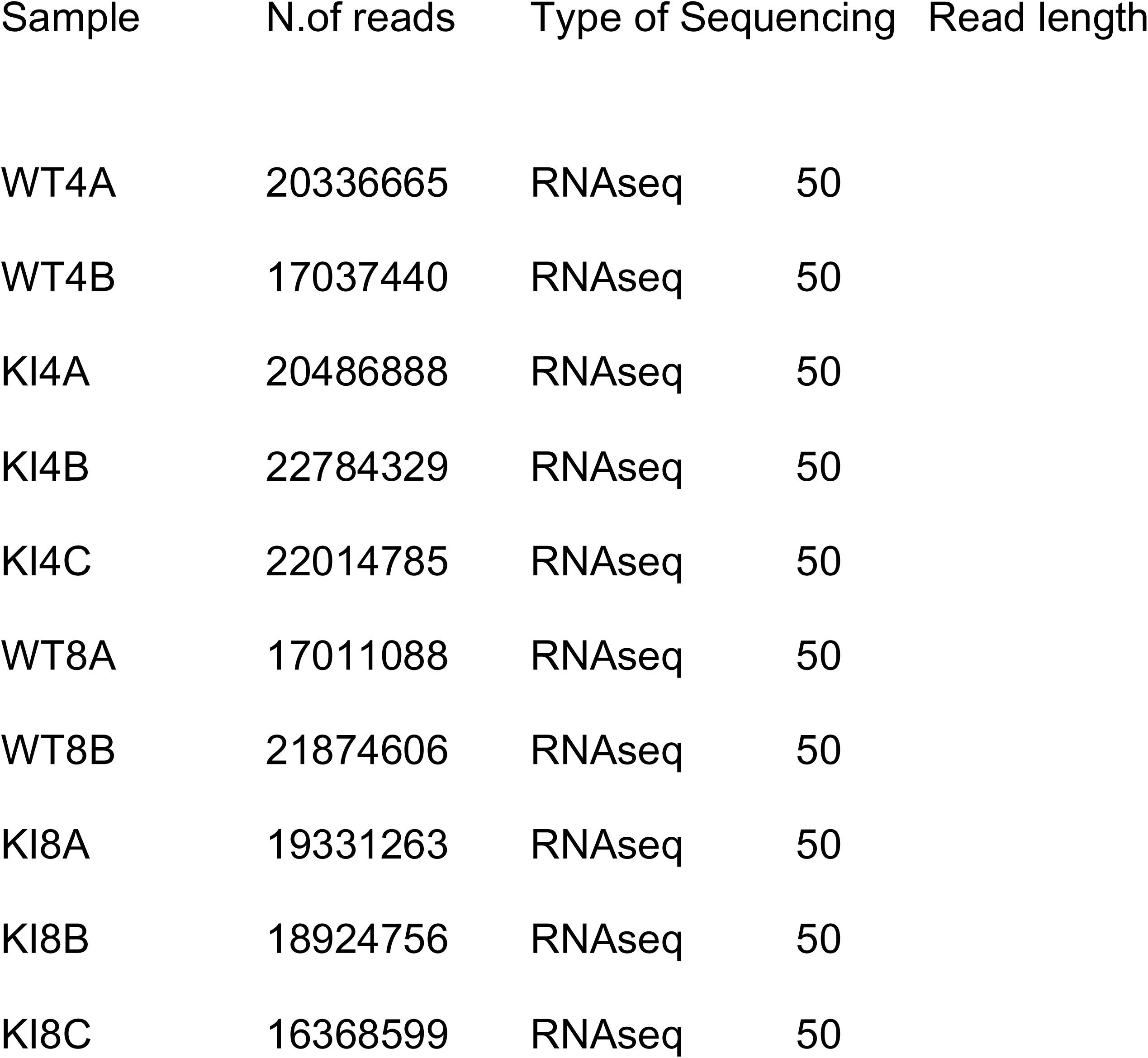

| Sample | N.of reads | Type of Sequencing | Read length |
| --- | --- | --- | --- |
| WT4A | 20336665 | RNAseq | 50 |
| WT4B | 17037440 | RNAseq | 50 |
| KI4A | 20486888 | RNAseq | 50 |
| KI4B | 22784329 | RNAseq | 50 |
| KI4C | 22014785 | RNAseq | 50 |
| WT8A | 17011088 | RNAseq | 50 |
| WT8B | 21874606 | RNAseq | 50 |
| KI8A | 19331263 | RNAseq | 50 |
| KI8B | 18924756 | RNAseq | 50 |
| KI8C | 16368599 | RNAseq | 50 |

Datasets can be accessed at the GEO repository (GSE193498).

Go to https://www.ncbi.nlm.nih._Enter token **ynirkwgshryzlkt** into the box

### In utero electroporation

In utero electroporation was performed as in Morenilla-Palao et al. (Morenilla-Palao et al., 2020) and sections of the SC of P11 electroporated mice were obtained and visualized as in (Benjumeda et al., 2013). pSilencer siRNA Expression Vector was purchased in Ambion. The Syt13 RNAi target sequence used to downregulate Syt13 was: 5’-GGCTGAGTTATTTGTGACA-3’.

### Analysis of axon terminals occupancy in the SC of electroporated mice

Analysis was performed using FiJi/ImageJ software (Schindelin et al., 2012). We selected the three most central 70-micron sections from the Superior Colliculus. In each section, a region of interest (ROI) was drawn to select and meassure the total area of the SC along the anterior-posterior axis using the DAPI channel as reference. For each ROI, the image was thresholded to detect GFP positive pixels. The percentage of the SC occupied by GFP positive axons was expressed as the ratio of the area occupied by GFP positive pixels by the total area of the SC. We call this value the “% GFP-covered Area”.

## DATA AVAILABILITY STATEMENT

The RNAseq datasets generated in this study are deposited at the NCBI Gene Expression Omnibus (GEO) with accession code GSE193498. All other included data in this study are available from the corresponding authors upon reasonable request. Analysis scripts are available on reasonable requests.

## ACKNOWLEDGMENTS

The laboratory of EH is funded with the following grants: PID2019-110535GB-I00 from the National Grant Research Program, PROMETEO Program (PROMETEO/2020/007) from Generalitat Valenciana, 20191956 from the Ramón Areces Foundation, an ERC-282329 grant from the European Research Council. The laboratory of AG is funded by PID2019-108194RB-100 from the National Grant Research Program. We also acknowledge the financial support received from the “Severo Ochoa” Program for Centers of Excellence in R&D (CEX2021-001165-S).

## STATEMENT OF COMPETING INTERESTS

All authors declare that they have no competing interests.

## AUTHOR CONTRIBUTIONS

SN performed and analyzed the CTB-tracing experiments. C M-P generated the Kir2.1 transgenic mice and extracted retinal RNA for RNA-seq experiments. She also recorded the retinal waves together with PO. SS analyzed the retinal waves. MH performed in situ hybridization and *in utero* electroporation experiments. MT L-C performed the computational analysis of RNA-seq data. DF perfomed electrophysiology experiments and AG analyzed the electrophysiology data. EH and SN wrote the original draft and EH designed, conceived and supervised the study.

## Notes

### Competing Interest Statement

The authors have declared no competing interest.

