## Supplementary figures and images for "Optimal frequency of perinatal retinal waves is essential for the precise wiring of visual axons in non-image forming nuclei"

### Suplementary Figure 1

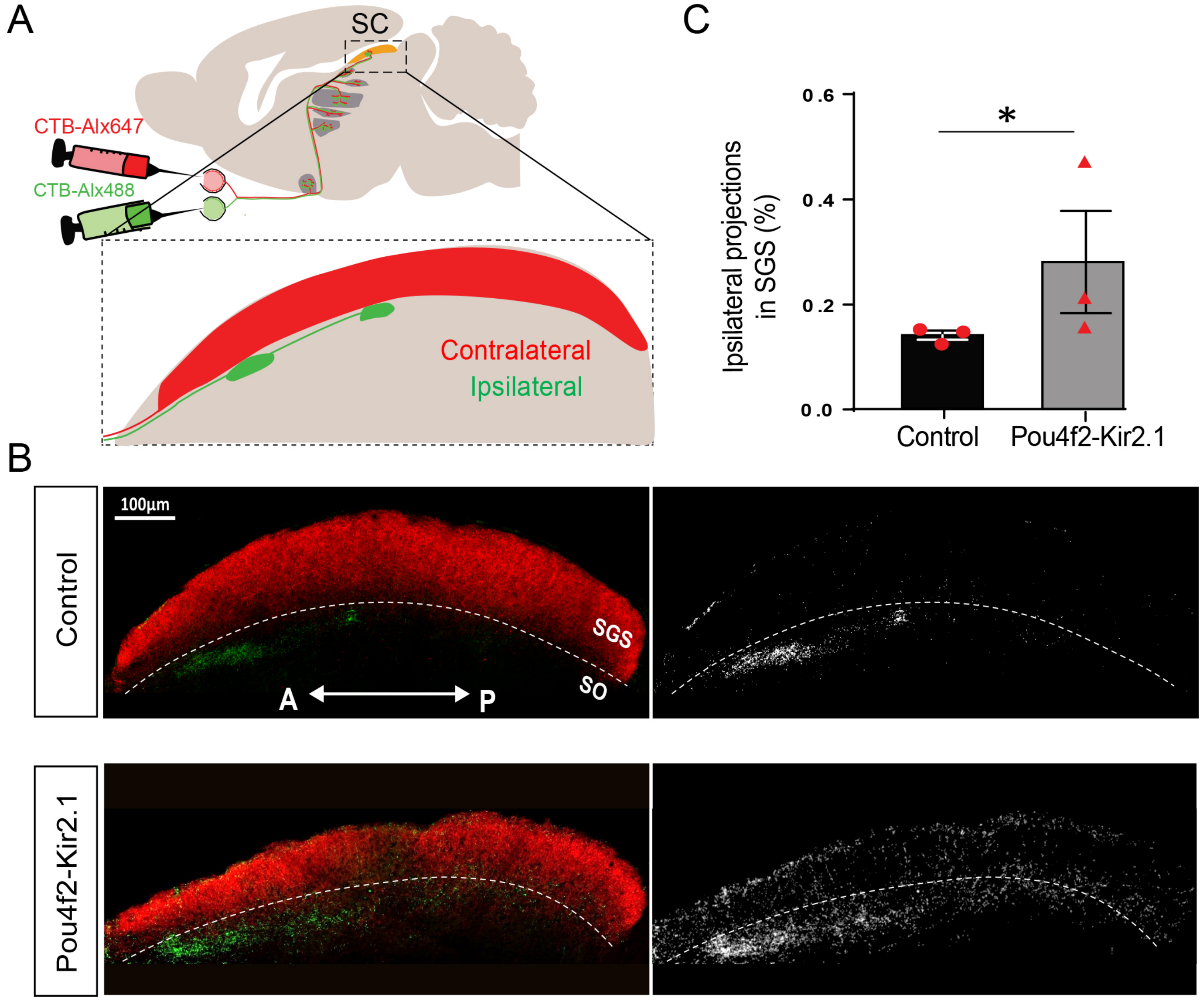

### Supplementary Figure 2

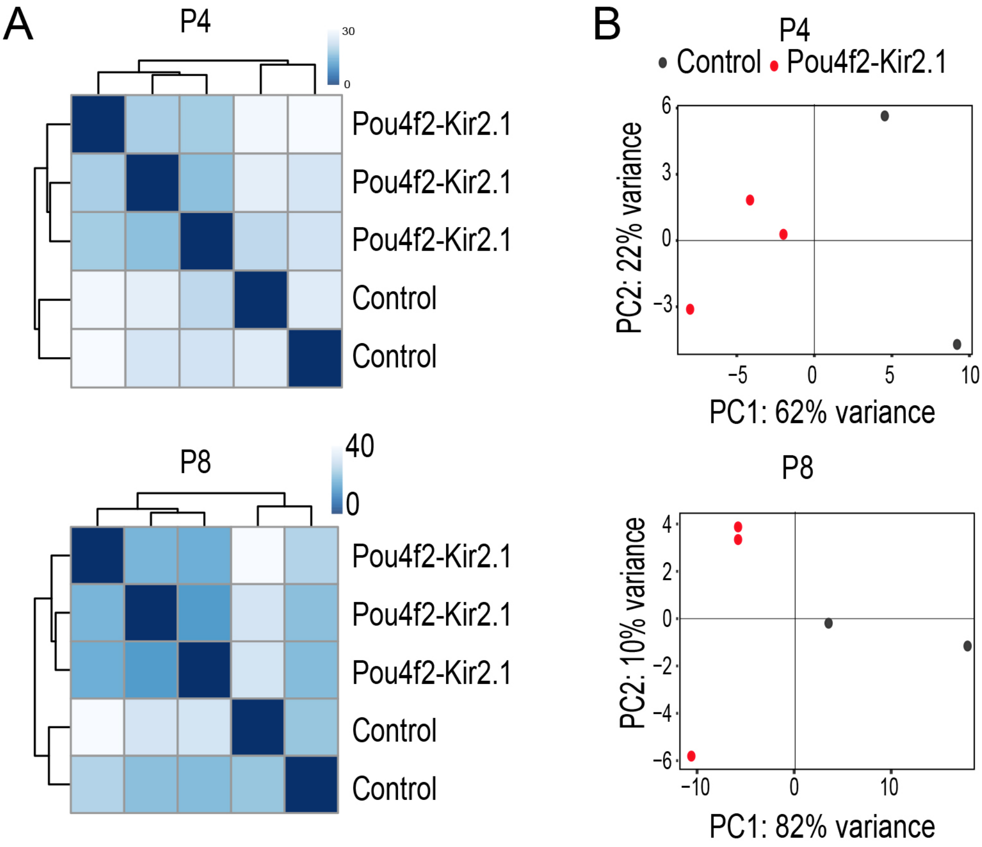

### Supplementary FIgure 3

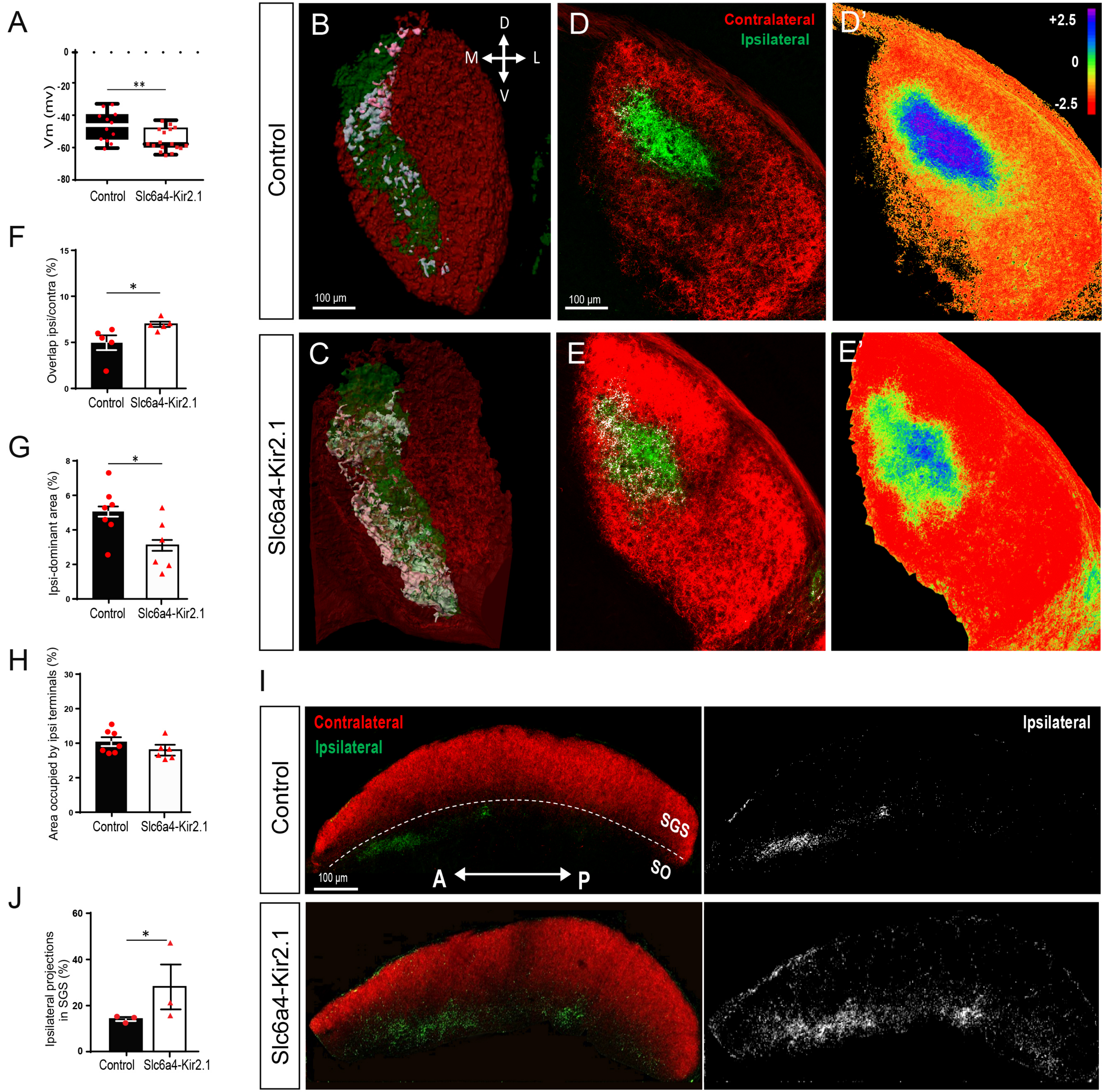
