## Supplementary material for "Optimal frequency of perinatal retinal waves is essential for the precise wiring of visual axons in non-image forming nuclei": Figure Legends to Suppl. Figures

**Supplementary Figure Legends**

**Figure S1 (Related to Figure 2). Pou4f2-Kir2.1 mice show eye specific segregation defects in the SC. (A)** Scheme illustrating the experimental approach consisting in intraocular injections of CTB fused to different Alexa fluorophores for each eye and analysis of coronal sections at the level of the SC (yellow). **(B)** Sagital sections from P11 Pou4f2-Kir2.1 and control mice show contralateral and ipsilateral projections at the level of the SC. Right panels show only the ipsilateral projections. **(C)** Quantification of the area occupied by ipsilateral terminals in the Stratum Griseum Superficialis (SGS). Each biological replicate (red dots) represents mean area values from three central sagittal sections of one SC/animal. Error bars indicate ±SEM (*$p < 0.05$, Student's unpaired T-test).

**Figure S2** (**Related to Figure 3). (A)** Pearson correlation matrix between normalized RNA-seq samples clustered by Euclidian dendrogram. **(B)** Principal component analysis of RNA-seq samples at P4 and P8.

**Figure S3 (Related to Figure 6). Slc6a4-Kir2.1 mice show impaired eye-specific segregation in the dLGN and SC. (A)** Averaged resting membrane potential in control cells and cells expressing Kir2.1. Dots represent TdTomato positive cells from retinas from three different mice. Whiskers extend to the min and max values (*$p < 0.05$, ** $p<0.01$, Student's unpaired T-test) **(B, C)** 3D reconstructions and **(D, E)** coronal sections through the dLGN of P11 Slc6a4-Kir2.1 mice and control littermate mice injected with CTB-Alexa647 and CTB-Alexa488. Overlaping of ipsilateral and contralateral terminals is visualized in white. **(D'-E')** R-distribution (log[ipsilateral/contralateral]) representing contralateral dominant area in red, unsegregated in green and ipsilateral dominant in blue. **(F)** Percentage of the area containing ipsilateral and contralateral terminals relative to the total area occupied by RGC terminals at the dLGN. **(G)** Percentage of the area with a majority of ipsilateral terminals relative to the total area occupied by RGC terminals. **(H)** Percentage of the area occupied by ipsilateral projections related to the total area occupied by RGC terminals. **(I)** Sagital sections from P11 mice at the level of the SC from Pou4f2-Kir2.1 and control mice show contralateral and ipsilateral projections. Ipsilateral projections are also shown in B&W at right panel. **(J)** Quantification of the area occupied by ipsilateral terminals in the Stratum Griseum Superficialis (SGS). Each biological replicate (red dots) represents mean values from three central coronal (dLGN) or sagittal (SC) sections from one nucleus/animal. Error bars indicate ±SEM (*$p < 0.05$, Student's unpaired T-test).

**Video 1.** Representative time-lapse of calcium waves in a flattened wholemount retina from a Pou4f2-;GCaMP6f P4 mouse. The video represents 2 min 30 s of recording at 1.8X speed. Scale bar: 300μm

**Video 2.** Representative time-lapse of calcium waves in a flattened wholemount retina from a Pou4f2-;Kir2.1YFP$^{flxStop}$;GCaMP6f P4 mouse. The video represents 2 min 30 s of recording at 1.8X speed. Scale bar: 300μm

**Table1 Contents.**

Sheet1. Index of the different tables

Sheet2. Differential expression analysis of WT vs mutants samples a P4 and P8)

Sheet3. Statistically significant DEGs (Padj <0,1)

Sheet4. Gene Ontology analysis of DEGs WT vs mutant samples at P4 and P8

Sheet5. Differential expression analysis of WT samples at P4 and P8 (Statistically significant DEGs (Padj <0,1)

Sheet6. Quality controls in the alignment of the raw data samples during the analysis of the data.
